# Modeling the radiative, thermal and chemical microenvironment of 3D scanned corals

**DOI:** 10.1101/2023.01.31.526450

**Authors:** Swathi Murthy, Cristian Picioreanu, Michael Kühl

## Abstract

1. Reef building corals are efficient biological collectors of solar radiation and consist of a thin stratified tissue layer spread over a light scattering calcium carbonate skeleton surface that together construct complex three dimensional (3D) colony structures forming the foundation of coral reefs. They exhibit a vast diversity of structural forms to maximize photosynthesis of their dinoflagellate endosymbionts (Symbiodiniaceae), while simultaneously minimizing photodamage. The symbiosis takes place in the presence of dynamic gradients of light, temperature and chemical species that are affected by the interaction of incident irradiance and water flow with the coral colony.
2. We developed a multiphysics modelling approach to simulate microscale spatial distribution of light, temperature and O_2_ in coral fragments with accurate morphology determined by 3D scanning techniques.
3. Model results compared well with spatial measurements of light, O_2_ and temperature under similar flow and light conditions. The model enabled us to infer the effect of coral morphology and light scattering in tissue and skeleton on the internal light environment experienced by the endosymbionts, as well as the combined contribution of light, water flow and ciliary movement on O_2_ and temperature distributions in the coral.
4. The multiphysics modeling approach is general enough to enable simulation of external and internal light, O_2_ and temperature microenvironments in 3D scanned coral species with varying degrees of branching and morphology under different environmental conditions. This approach is also relevant for simulating structure-function relationships in other benthic systems such as photosynthetic biofilms and aquatic plant tissue, and can also be adapted to other sessile organisms such as symbiont-bearing giant clams, ascidians, jellyfish or foraminifera. The model could also be useful in more applied research such as optimization of 3D bioprinted constructs where different designs can be evaluated and optimized.

## 1. Introduction

Reef building, scleractinic corals construct a complex three dimensional (3D) calcium carbonate skeleton, which is the framework for coral reef ecosystems of considerable biological and socio-economic importance (Costanza et al., 2014). Tropical, scleractinic corals rely on endosymbiotic dinoflagellate algae from the family Symbiodiniaceae, which are hosted in the coral endoderm tissue and excrete photosynthates to the host, providing the majority (up to >90%) of carbon needed for coral animal respiration along with O_2_ (Muscatine, 1973). Symbiont photosynthesis can also enhance coral calcification (Chalker, 1981; Goreau et al., 1996). In exchange, the host provides the endosymbionts with a conducive and protected environment and a steady, albeit limited supply of nutrients.

However, the coral-algal symbiosis can rapidly deteriorate under environmental stress (LaJeunesse et al., 2018), (Moberg & Folke, 1999). Environmental factors related to ongoing climate change such as ocean acidification (Jackson et al., 2001; van der Zande et al., 2020), deoxygenation (Hughes et al., 2020) (Altieri et al., 2017) and warming (Hughes et al., 2017), increasingly result in severe mass bleaching and mortality of corals (Knowlton et al., 2021). Coral bleaching and mortality varies as a consequence of solar irradiation levels (Dunne & Brown, 2001; Mumby et al., 2001), local seawater temperature variation (Teneva et al., 2012), coral colony morphology (Loya et al., 2001; Van Woesik et al., 2012), water flow around the coral (Jimenez et al., 2008; Nakamura et al., 2003), and O_2_ levels in the surrounding water (Altieri et al., 2017; Hughes et al., 2020; Johnson et al., 2018). Hence, it is important to improve the mechanistic understanding of how the photobiology and physiological activity of the coral holobiont are affected by macroscale colony morphology and microscale variations in tissue/skeleton architecture, and how interactions with the incident solar irradiance and flowing seawater modulate the physico-chemical microenvironment and metabolic activity of corals. This will enable us to better estimate hotspots for coral bleaching, the coral reef’s response to environmental stressors, and the relationship between the host microenvironment and microbiome/endosymbiont populations. Understanding of light propagation, heat and mass transfer in relation to the 3D morphology of a coral colony can also provide insight into the mechanisms governing spatial and temporal variability in coral bleaching and recovery within and between coral species.

Corals are efficient biological collectors of solar radiation (D. Wangpraseurt et al., 2014) and consist of a thin (from ~100 μm to several mm) stratified tissue layer spread over a light-scattering skeleton matrix (Davy et al., 2012). They are exposed to a wide range of light environments ranging from super-saturating, broadband (UV to near infrared) solar irradiance in the shallow reefs (Jimenez et al., 2012; Veal et al., 2010; Wangpraseurt et al., 2014) to very low illumination in the blue-green spectral range at mesophotic depths (Eyal et al., 2015; Tamir et al., 2019). The morphology of the colony and the surrounding reef, along with the optical properties of the host tissue and skeleton, can lead to variable light conditions experienced by different parts of a coral colony and hence the endosymbionts (Kaniewska et al., 2011; Wangpraseurt et al., 2014).

Corals have developed a vast diversity of structural forms and mechanisms at cellular, polyp and colony level, to maximize photosynthesis of their endosymbionts under spatial and temporal variation in light availability (Anthony & Hoegh-Guldberg, 2003; Kaniewska et al., 2011; Todd et al., 2008), while simultaneously minimizing photo-damage (Brodersen et al., 2014; Terán et al., 2010) and oxidative stress (Pacherres et al., 2022), and maximizing solute and heat exchange (Jimenez et al., 2011). Consequently, corals can reach high photosynthetic quantum efficiencies close to 0.1 O_2_ photon^-1^, approaching theoretical limits, under moderate irradiance levels (Brodersen et al., 2014; Dubinsky et al., 1984).

Light absorption by corals not only drives symbiont photosynthesis *in hospite*, but can also affect the thermal microenvironment via local heating (Jimenez et al., 2008), as most of the absorbed light energy is dissipated as heat (Brodersen et al., 2014). Light is mainly absorbed in the coral tissue and, to some extent, in the skeleton by endolithic algae and photosynthetic bacteria (Kühl et al., 2008). However, both coral morphology (Kaniewska et al., 2011; Kaniewska & Sampayo, 2022; Kramer et al., 2022) and the inherent scattering properties of coral skeleton and tissue modulate light propagation and thus the light exposure of the microalgal symbionts (Enríquez et al., 2005; Wangpraseurt et al., 2016) (Bollati et al., 2022; Lyndby et al., 2016).

The morphology and the degree of branching in coral colonies affect not only the light field but also the water flow around the colonies. The flow determines the thickness and shape of boundary layers for momentum transfer (MBL, i.e., water velocity change near the coral surface), mass transfer (DBL, i.e., concentration change in the diffusion-dominated region adjacent to the coral tissue) (Chan et al., 2016), as well as heat transfer (TBL, referring to temperature changes) (Jimenez et al., 2008; Ong et al., 2019). The coral consumes part of the O_2_ produced by the symbionts under light (Al-Horani et al., 2003), while an eventual surplus of oxygen and heat generated in the tissue is transported into the surrounding water column and the coral skeleton. By introducing a resistance to mass and heat transfer, boundary layers decrease the mass and heat exchange rates between the bulk water and the coral, to the extent in which solute diffusion (Shashar et al., 1993) and heat conduction (Ong et al., 2012; Ong et al., 2017; Ong et al., 2019) can become bottlenecks. These transfer resistances can be alleviated to some extent by actively decreasing boundary layer thickness through an intensified flow, as achieved by tissue and polyp level changes like contraction-expansion and ciliary movement that can enhance mass (Pacherres et al., 2020; Pacherres, 2022; Shapiro et al., 2014) and heat (Ong et al., 2017) transfer, along with affecting light penetration (Wangpraseurt et al., 2014; Wangpraseurt et al., 2017). The resulting spatial and temporal variations in temperature distribution and concentration of chemical species (e.g., dissolved oxygen, inorganic carbon and pH) affect the coral microenvironment and can form distinct microhabitats within a given coral colony. Such fine scale ecological niche heterogeneity may affect the composition of both endosymbionts and microbiomes across the different compartments of the coral holobiont, but the microscale fitness landscape of corals remains largely unexplored, in part due to technical challenges (Hughes et al., 2022).

Numerical modelling is a powerful way to integrate the physical, chemical and biological complexity (at several spatial and temporal scales) into a systematic framework, with the aim to describe, understand and ultimately predict coral responses to a changing environment. Modelling the growth of coral colonies in response to environmental parameters has been pioneered by the work of Kaandorp and coworkers (Chindapol et al., 2013; Kaandorp, 2013). Several studies have also modeled the interaction of light and fluid flow with colony morphology and the resulting surface temperature (Ong et al., 2012; Ong et al., 2018; Ong et al., 2017; Ong et al., 2019) and mass transfer at the coral surface (Chang et al., 2014; Shapiro et al., 2014). However, these models have not included radiative transfer in the tissue-skeleton matrix, or simulations of internal gradient of light, temperature and chemical parameters across the tissue and skeleton.

Empirical and theoretical models have shown how skeleton-dependent scattering can enhance the local light field and absorption of Symbiodiniaceae *in hospite* (Enríquez et al., 2005; Swain et al., 2016). Ray tracing (Ong et al., 2018) and probabilistic Monte Carlo (MC) modeling techniques for light propagation (Terán et al., 2010; Wangpraseurt et al., 2016) assuming a simple two-layer tissue-skeleton geometry have also been employed. Recent advances in estimating inherent optical properties of different species of living corals (Jacques et al., 2019; Spicer et al., 2019; Wangpraseurt et al., 2018) now enable more accurate simulation of internal light fields in specific corals.

We recently developed a numerical model of a stratified coral tissue on top of skeleton to link internal light distribution to light absorption, radiative heat dissipation, heat transfer and mass transfer of photosynthetically produced O_2_ (Taylor Parkins et al., 2021). The model couples a Monte Carlo (MC) simulation of light propagation with numerical modelling of heat production and metabolism inside the coral to simulate irradiance, temperature and O_2_ microprofiles at small scale in simplified, schematic geometries. In the present study, we have expanded this multiphysics modelling approach to simulate microscale distribution of light, temperature and O_2_ in and around branched coral fragments (cm scale) with complex, natural morphology, as determined by 3D scanning techniques. We validate the model by comparing simulated microenvironmental parameters over the coral topography with the corresponding microscale measurements on the same fragment under similar irradiance and laminar flow conditions. This enabled us to describe the effect of coral morphology on the internal light environment and the combined effects of light, water flow and ciliary movement on the O_2_ and temperature distribution in the coral. The presented modeling approach can easily be adapted for simulating effects of flow and irradiance on the optical, thermal and chemical microenvironment in other types of aquatic organisms and systems with a defined structural composition, either reflecting their natural structure, as obtained from 3D scanning, or more schematic idealizations, e.g. in bionic 3D bioprinted constructs.

## 2. Materials and methods

The methodological approach involved an experimental and a theoretical part. First microsensor measurements of light, O_2_ and temperature were performed on a fragment of the branched coral *Stylophora pistillata* (Figure 1a,b) in a flow chamber under defined irradiance and flow conditions. The fragment was subsequently 3D scanned and meshed (Figure 1c,d), followed by multiphysics 3D modeling of the radiative, heat and mass transfer of the meshed structure (Figure 1f) and subsequent comparison of simulated distributions of light, O_2_ and temperature with the experimental measurements (Figure 1e).

**Figure 1:**
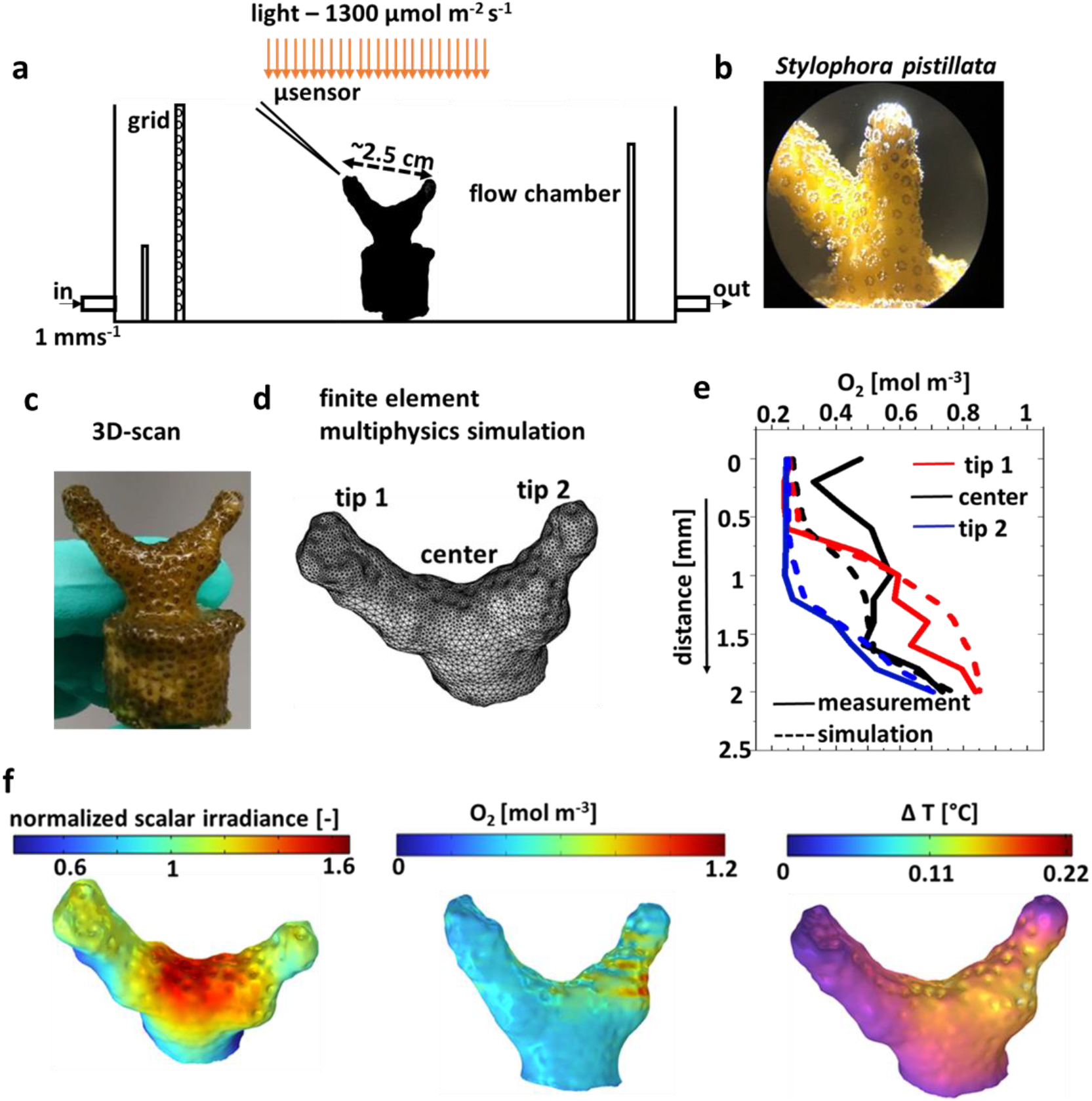
Overview of experimental and modeling approach. Experimental flow chamber set-up for microsensor measurements (a) on a coral fragment of *Stylophora pistilla* (b,c). Tetrahedral mesh based on the 3D scan of the coral (d), used for multi-physics simulation. Comparison of microsensor measurements and simulations (here illustrated with measured and simulated O_2_ concentration profiles) in particular areas of the coral (e). Simulated surface values of normalized scalar irradiance, O_2_ concentration and temperature at the coral tissue-water interface (f).

### 2.1 Coral specimens and husbandry

Colonies of the branched coral *Stylophora pistillata* were obtained from a sustainable, commercial provider (Dejong Marine Life, Netherlands). Colonies were fragmented into smaller nubbins that were mounted in small plastic caps using AF Gel Fix (https://aquaforest.eu) and kept in dedicated aquaria supplied with continuous flow of artificial reef water at 25-26°C, salinity of 36 g/kg, and moderate levels of downwelling photon irradiance (150 - 200 μmol photons m^-2^ s^-1^; 400–700 nm) provided over a 12:12 hour dark-light cycle by a programmable aquarium lamp (MAXSPECT R5X 150W; Decocean Aquarium).

### 2.2 Experimental setup

A small fragment of *S. pistillata* (Figure 1b) was placed in a custom-made black, acrylic flow chamber, and measurements were done with two different orientations of the fragment relative to the laminar flow. The samples were placed for at least 1 hour in the chamber prior to microsensor measurements to ensure steady-state conditions. The coral fragment was continuously flushed with aerated seawater at 26°C and salinity of 35 g/kg. An average flow velocity of ~1 mm s^-1^ was maintained by a water pump connected to the flow chamber and submerged in the thermostated aquarium reservoir.

The sample was illuminated with a constant downwelling photon irradiance of 1300 μmol photons m^-2^ s^-1^ from a fibre-optic tungsten halogen lamp equipped with a heat filter and a collimating lens (KL-2500LCD, Schott GmbH, Germany), positioned vertically above the flow chamber (Figure 1a). The downwelling photon irradiance (Ed in units of μmol photons m^-2^ s^-1^) of photosynthetically active radiation (PAR; 400–700 nm) was measured with a calibrated photon irradiance meter (ULM-500, Walz GmbH, Germany) equipped with a planar cosine collector (LI-192S, LiCor, USA) positioned in the light path approximately at the same distance as the coral.

### 2.3 Microsensor measurements

Measurements were conducted on the connective tissue, i.e., coenosarc, between individual polyps, which had a more even topography and less contractile tissue as compared to polyp tissue.

#### Light measurements

Spectral scalar irradiance was measured using a fibre-optic scalar irradiance microprobe with a spherical tip diameter of ~100 μm (Rickelt et al., 2016) connected to a fibre-optic spectrometer (USB 2000+, Ocean Optics, USA). The incident downwelling irradiance was measured at the same height as the coral surface by placing the light sensor in a black (non-reflective) light well under the vertically incident light in the flow chamber. All spectral scalar irradiance measurements at the coral tissue surface were normalized to the incident spectral irradiance. The results are represented as mean ± standard deviation, averaged over 12 measurement points taken close to each other in a given region of interest (as indicated by squares in Figure 2c) and for the 2 different orientations of the fragment with respect to the flow.

**Figure 2:**
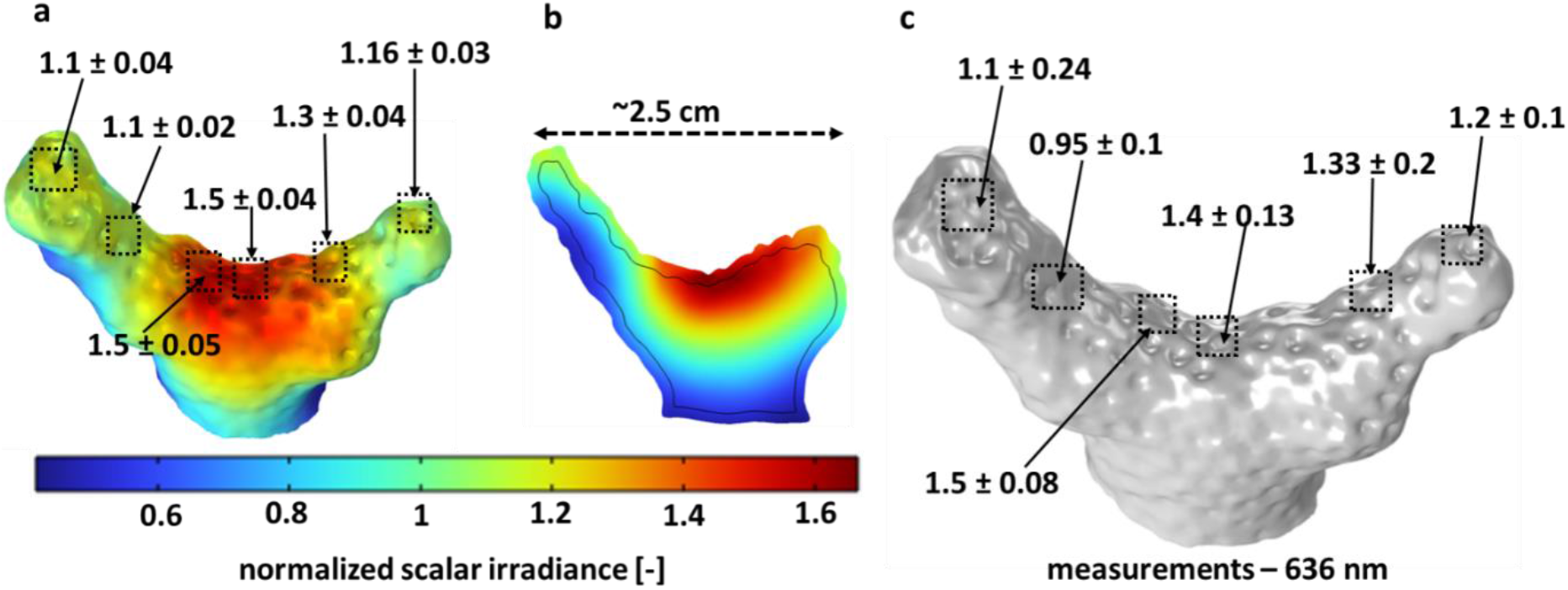
Comparison between measured and simulated scalar irradiance (636 nm; ratio of scalar irradiance to incident irradiance). (a) Simulated scalar irradiance distribution over the coral tissue-water interface; (b) Cross-section of simulated scalar irradiance depth distribution along the middle section of the fragment; (c) Measured scalar irradiance in similar areas, from which simulated values were extracted. Optical properties (OP1) used for simulation are listed in Table S1. The values shown here represent means ± standard deviation, averaged over 12 different points in each area of interest as indicated by the squares.

#### Oxygen measurements

Oxygen concentrations were measured with Clark-type microelectrodes (tip diameter approx. 25 μm; OX-25, Unisense A/S, Aarhus, Denmark)(Revsbech, 1989). The microsensor was connected to a pA-meter (Oxymeter, Unisense A/S) interfaced to a PC. The O_2_ microsensors were linearly calibrated from sensor signal measurements in aerated and anoxic seawater at experimental temperature and salinity.

#### Temperature measurements

Thermocouple microsensors (tip diameter approx. 50 mm; T50, Unisense A/S) were connected to a thermocouple meter (Unisense A/S). A high precision thermometer (Testo 110, Testo AG, Germany) was used to linearly calibrate the temperature microsensor signal from readings in seawater at different temperatures.

For measurements, the microsensors were mounted on a motorized micromanipulator (MU-1, PyroScience, GmbH), which was interfaced to a PC and controlled by dedicated data acquisition software (ProFix, Pyros-Science GmbH). All measurements were made at an angle of 45° relative to the vertically incident light beam to avoid shading. The measurements were made at the coenosarc tissue in three different regions of the coral fragment as shown in Figure 1d. The temperature profiles were represented as the temperature increase relative to the water temperature (in the free flowing region) directly above the measurement point. The O_2_ and temperature profiles were measured from the water column into the coenosarc in vertical steps of 100 μm, as described previously (Jimenez et al., 2008). Scalar irradiance measurements were done by placing the sensor tip at the coral tissue surface. Positioning of microsensor tips relative to the coral tissue surface was monitored visually by observation under a dissection scope.

### 2.4 3D scanning and OCT imaging

We used a PC-controlled, structured light 3D scanner (Einscan-SP, Shining 3D) to generate non-texture 3D scans of coral fragments in air. A sample was placed on the turn table of the calibrated scanner. Data acquisition and processing was controlled by the system software (EXScan S_V3; Shining 3D). The brightness setting of the scanner was adjusted to avoid saturation of the camera signal. A 360° scan of the fragment was made under different orientations, to capture the complete morphology of the fragment avoiding missing faces. The different scans were aligned using the “align by feature” mode in the scanner software. A watertight mesh of the model was exported as an STL file. The fragment was scanned with and without the tissue layer to estimate an average tissue thickness, from the volume difference between the two scans. The tissue was removed by immersing the fragment in 30% hydrogen peroxide solution overnight. This procedure led to a tissue thickness estimation of ~900 μm. However, this could be an overestimate as the reconstruction software fills in holes in the scan by extrapolation, in places where it could not scan the object correctly (e.g., due to complex shape and highly absorbing or reflective surfaces).

For an alternative determination of the coral tissue thickness, we used a 930 nm spectral domain OCT system (Ganymed II, Thorlabs, Germany) equipped with an objective lens with an effective focal length of 18 mm and a working distance of 7.5 mm (LSM02-BB; Thorlabs GmbH, Dachau, Germany) for OCT imaging of corals immersed in seawater with a maximal axial and lateral resolution in water of 5.8 μm and 8 μm, respectively (Wangpraseurt et al., 2017). Two-dimensional OCT B-scans were acquired at a fixed pixel size of 581 x 1024. The scans were used to estimate the tissue thickness, however, due to shadowing effects and uncertainties in the exact refraction index of the coral tissue, the OCT measurements underestimated tissue thickness. For the simulations, we therefore selected a tissue thickness of 650 μm, which was inbetween the values from the OCT and 3D scans.

### 2.5 Numerical modeling

A three-dimensional mathematical model was constructed with the aim of simulating the spatial distribution of dissolved oxygen and temperature around and within the coral, as influenced by the light transport and by the water flow. Model predictions were compared to the measured profiles of light, O_2_ and temperature on the coral fragment.

For the simulations, the scanned 3D coral geometry was assumed to consist of a tissue layer with an uniform thickness created on top of the skeleton (Figure S1a, S1b), which was placed in a rectangular flow chamber (Figure S1c). Different sub-layers within the coral tissue (Taylor Parkins et al., 2021), with varying material properties and supporting different chemical reactions, were represented indirectly as function of the distance from the tissue-water surface. The model first computes the light field within and around the coral (Figure S1d), then the laminar flow of water around the coral in the flow chamber. Subsequently, mass and heat balances allow calculation of the radiation driven O_2_ and temperature distribution in the stratified coral domain and the water column (Figure S1e and S1f). All the model parameters are listed in Table S1 (supplementary information).

#### Model geometry

The model geometry consisted of a solid domain, the 3D scan of the coral fragment, enclosed by the liquid in a box matching the dimensions of the flow chamber used experimentally (Figure S1c). This geometry was built in COMSOL Multiphysics (v6, COMSOL Inc., Burlington, MA) based on the STL file provided by the coral 3D scanning. Two different orientations of the fragment with respect to the flow (Figure S1) were simulated.

An essential part of the model includes the effect of coral tissue sub-layers with various properties, thus assigning different material properties and reactions as a function of depth within the tissue, as done in (Taylor Parkins et al., 2021) albeit in a much simpler geometry. However, an explicit partitioning of the large 3D coral domain in several very thin subdomains on the skeleton surface proved to be very difficult computationally, especially regarding the accurate meshing of these thin layers. We therefore adopted the solution of representing these layers implicitly, by a distance *d* from the coral/water surface. The perpendicular distance (*d*) within the coral from the coral/water surface was computed in COMSOL (the wall-distance interface) by a modified Eikonal equation (Fares & Schröder, 2002). Consequently, the optical, mass and heat transfer properties characteristic for different tissue sub-layers were defined as a function of this distance, *d*.

#### Radiative transfer

The scalar irradiance at different positions in the coral and water was determined by calculating the radiative transfer (Figure S1d) using the 3D Monte Carlo (MC) approach implemented in the free software ValoMC (Leino et al., 2019), as described in our previous work (Taylor Parkins et al., 2021). The simulation of photon transport was performed by launching photons with a wavelength of 636 nm, within the Chl *c* and Chl *a* absorption band of the coral microalgal symbionts. The present model ignores fluorescence and takes only scattering and absorption into account. Various tissue layers and skeleton optical properties (OP1, Table S1) were assigned as a function of the calculated wall distance *d*.

The simulated point cloud of scalar irradiance values from ValoMC were imported into COMSOL and mapped over the 3D geometry by solving Poisson’s diffusion equation in weak form, in order to smoothen the variations in the scalar irradiance data inherent due to the stochastic nature of the MC simulation:

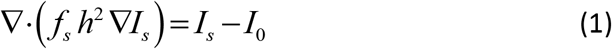

where *I_s_* is the smoothed normalised scalar irradiance, *I_0_* is the initial normalised scalar irradiance, *h* is the local mesh element size, and *f_s_* is a smoothing factor (0.1; 0 < *f_s_* < 1). A zero light flux condition was applied to all boundaries.

The simulated scalar irradiance at different positions in the geometry was normalized to the incident scalar irradiance. The results are represented as mean ± standard deviation, averaged over 12 different points, sampled close to each other, as indicated by the squares in Figure 2a.

#### Fluid flow

Stationary incompressible Navier-Stokes equations for laminar flow (Reynolds number of ~9) were used to simulate the water flow around the coral fragment using COMSOL:

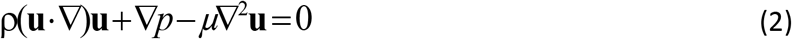

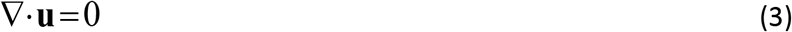

where **u** is the velocity vector, *p* the pressure, *μ* the dynamic viscosity and *ρ* the density of water. The water inflow had an average velocity of 1 mm s^-1^, while a zero gauge pressure was set in the outflow. Top, bottom and lateral walls of the flow cell were no-slip (zero-velocity). Two cases were assumed for flow at the coral surface: zero-velocity (without ciliary movement) and a set velocity (with ciliary movement). Ciliary movement as proposed in other studies (Pacherres et al., 2020; Shapiro et al., 2014) was included as cilia-induced currents by assuming an oscillating horizontal velocity component at the coral surface as (Pacherres et al., 2022):

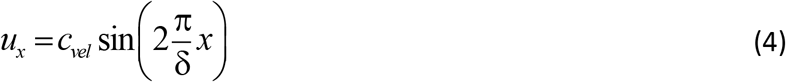

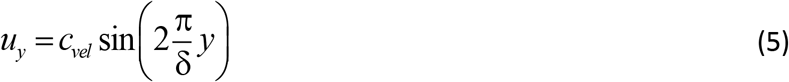

with *x* and *y* being the distances from surface, *C_vel_* the maximum ciliate beating velocity taken 150 μm s^-1^ as in the previous measurements (Shapiro et al., 2014) and δ the characteristic length scale of the vortices (assumed 1200 μm, close to the average calyx size of the coral).

The simulated flow profile around the coral was similar with and without the base onto which the coral was glued (Figure S2). Hence, in order to simplify the geometry and the computational burden, all the simulations presented here were made without the base.

#### Oxygen transport and reactions

The dissolved oxygen concentration, *c_O2_*, in the coral and surrounding water was computed from a stationary material balance, written in a general form as eq. (6):

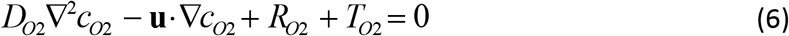

Diffusive transport was applicable in both domains with diffusion coefficient *D_O2_* taking the value *D_O2,w_* for dissolved O_2_ in water, but reduced values in the coral (0.5·*D_O2,w_* in the tissue and 0.01·*D_O2,w_* in the skeleton). In the water domain there were no reactions involving oxygen (*R_O2_*=0), while in the coral tissue/skeleton there was no convective transport (**u**=0). The gas-liquid transfer term *T_O2_* accounted for eventual O_2_ super-saturation, thus preventing apparition of exaggeratedly high O_2_ concentrations in certain areas. A constant O_2_ concentration *c_0,O2_* was imposed in the water inflow and the classical convection-only condition was assumed in the water outlet, with the rest of flow cell walls insulated (no flux of O_2_). The flux and concentration continuity were assumed on the coral/water interface.

The net photosynthetic O_2_ production rates in oral and aboral gastrodermis (*gas*) were calculated as a function of scalar irradiance *I_p_* using the average light absorption coefficient of the coral tissue μ*_a,tis_*:

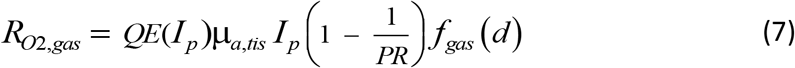

Here, the quantum efficiency of symbiont photosynthesis (mol O_2_ produced per mol photon) is represented as a function *QE*(*I_p_*) decreasing with increasing scalar irradiance (Figure S3, (Brodersen et al., 2014)). The rate is furthermore limited by the ratio of photosynthesis to respiration rate, *PR*=3.5 (Cooper et al., 2011). *f*_gas_ is a switch function depending on the distance *d* from the coral/water interface, which allows defining *R_O2_* only in the gastrodermis, i.e., *f*_gas_ = 1 if the distance is 270 – 370 μm for oral, 470 – 570 μm for aboral gastrodermis layer, and *f*_gas_ = 0 elsewhere.

The respiration rate in the epidermis layer was calculated as function of O_2_ concentration *c_O2_*:

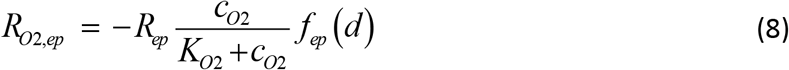

with *R_ep_* the coral host maximum respiration rate and *K_O2_* the half-saturation coefficient for O_2_ limitation for respiration. Again, the switch function *f_ep_* (*d*) activates this rate (*f_ep_* = 1) only at a distance *d* between 90 and 170 μm from the coral surface.

The O_2_ consumption in the coral skeleton by endolithic bacteria follows a similar rate:

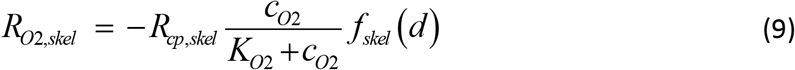

with *R_cp,skel_* the maximum O_2_ consumption rate and the same half-saturation coefficient *K_O2_*. *f_skel_* switches on this rate for *d* > 650 μm and off elsewhere.

Finally, the transfer of dissolved oxygen to gas bubbles at locations with O_2_ super saturation in water (at atmospheric saturation) and gastro-vascular cavity domains was included as:

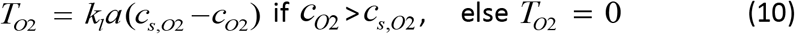

where *k_1_a* is a gas-liquid exchange coefficient and *c*_s,O2_ is the O_2_ solubility in water.

#### Heat generation and transport

The spatial temperature *T* distribution in coral and surrounding water was computed from the general heat balance equation including conduction, convection and source terms:

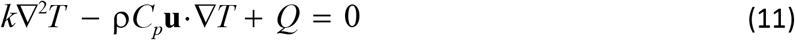

where *k* is the thermal conductivity of the specific layer, *C_p_* and ρ are the specific heat capacity and density of water, *Q* is the heat source and **u** is the vector of water velocity. In particular, heat conduction and generation terms were considered in the coral fragment, while conduction and convection were included in the water domain (neglecting heat generated from light absorption in water). The heat source, *Q*, originates from absorbed light only, which is proportional with the scalar irradiance, *I_p_*, and the light absorption coefficient, μ_*a*_, of the specific layer:

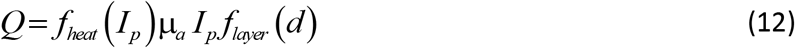

where switch functions *f_layer_* correspond to specific layers at different distanced from the coral/water surface. *f_heat_*(*I_p_*) is the fraction of light dissipated as heat in the oral and aboral gastrodermis (Figure S4, (Brodersen et al., 2014)), where *f_heat_*<1 because part of the light energy is used in photosynthesis, while *f_heat_* was set to 1 in the other tissue layers and skeleton.

The inflow of water had a temperature of *T_0_*, while a zero-temperature-gradient condition was set for the water outflow. Thermal insulation was set at all other flow-cell walls, while heat flux continuity was assumed at the coral/water interface. The thermal properties of the skeleton were taken from Jimenez et al (Jimenez et al., 2008), while the tissue thermal properties were assigned values similar to human tissue (Hasgall PA, 2018) (due to lack of other measurements).

### 2.6 Comparison between simulated and measured data

For comparison of measured and simulated scalar irradiance values, scalar irradiance microsensor measurements at 636 nm (normalized to incident irradiance) at the coral tissue/water interface were averaged in different areas of interest over the coral fragment, and were then compared to simulated scalar irradiance values in the same areas of interest (Figure 2). For comparison of measured and simulated O_2_ and temperature measurements, we compared measured profiles in particular positions over the coral structure with extracted simulated vertical profiles of O_2_ concentration and temperature over the same regions (Figure S5).

## 3. Results and discussion

We used a combination of microsensor measurements, 3D scanning and numerical modeling to investigate the influence of coral structure and morphology on the light, O_2_ and temperature microenvironment in and around a coral fragment (Figure 1). Microsensor measurements are local, making it unfeasibly tedious to map the entire sample and account for spatial heterogeneity and hot spots. Hence supplementing them with simulations of the light, O_2_ concentration and temperature distribution over the 3D morphology of actual samples can help identifying such areas of interest and lead to better data interpretation. On the other hand, supporting simulations with measurements is imperative to realize a physical meaning for the model output, which can also help fine tune certain assumptions, to achieve a more realistic output from the model. Such combination of simulations and microsensor measurements provides a powerful toolset for exploring the coral microenvironment under different flow and light scenarios.

### 3.1 Light microenvironment

The 3D light simulation generates the distribution of the scalar irradiance within the whole computational domain including water, coral tissue and skeleton. Computed scalar irradiance is displayed at the surface (Figure 2a) and within the coral fragment (Figure 2b). Both the simulations and the light measurements demonstrated the presence of a heterogeneous light field over and within the coral fragment. The computed scalar irradiance shows a fair match with the measurements (Figure 2c) at the same wavelength (636 nm).

Both simulations and the measurements showed that the surface scalar irradiance values (quantified as the ratio of the scalar irradiance over the incident irradiance; mean ± standard deviation, n=12 in each area of interest) were highest at the center of the fragment (measured: 1.40 ± 0.13; simulated: 1.50 ± 0.04; Figure 2a and 2c), where the tissue surface was relatively flat. At the tips of the fragment, the relative scalar irradiance values were lower because the surface was more inclined with respect to the incoming light.

The rear (shaded) sides of the fragment exhibited much lower scalar irradiance values around half of the incident irradiance (measured and simulated). The simulations indicate that the highest scalar irradiance was close to the tissue-skeleton interface in the middle of the fragment, reaching up to ~1.7 times the incident irradiance (Figure 2b).

Overall, the simulated scalar irradiance at the coral surface compared well to actual measured values in the same regions of the investigated coral. But we note that the light simulations relied on literature values of the inherent optical parameters of coral tissue and skeleton (Jacques et al., 2019), while the inherent optical properties of coral tissue and skeleton are more complex and involve e.g. different tissue layers with different refractive index (e.g. Wangpraseurt et al., 2014) and tissue regions with more or less scattering and absorption (Wangpraseurt et al., 2019). To illustrate how the model responds to differences in optical parameters, we also simulated the light field in *S. pistillata* fragments using optical parameters determined from OCT measurements (Wangpraseurt et al., 2019) (Figure S6, S7). While the absolute values of scalar irradiance enhancement changed between simulations with different optical parameters, the overall light distribution remained more or less identical between the simulations and reflected the pattern in the experimental light measurements. This highlights the fact that we are still lacking a thorough fine scale characterization of inherent optical properties of coral tissue and skeleton.

The light simulation and measurements revealed an interplay between skeleton and tissue optics that may be important in enhancing coral light harvesting. The light scattering from the skeleton leads to light enhancement in the tissue and a distribution of incident light to shaded areas. For a given direction of the incident light, the distribution of light can vary significantly, even across a small region of a coral colony, leading to hotspots and shaded regions. Light enhancement in a coral fragment is due to a combination of surface morphology and scattering in the tissue and skeleton, which enhances and redistributes light within the fragment (Wangpraseurt et al., 2016). Regions of a coral with relatively flat surfaces (angle of incident light relative to the tissue surface close to 0°) will experience less loss of light due to surface reflections arising from refractive index mismatch, as compared to more inclined surface areas in the colony experiencing lower incident irradiance (due to the cosine dependence of Fresnel reflection on the angle of incidence).

However, even shaded regions in a coral colony can receive some light due to lateral distribution of light via the tissue and skeleton (Enríquez et al., 2017; Wangpraseurt et al., 2016; Wangpraseurt et al., 2014).

The light field in corals is affected by the optical properties of tissue and skeleton, as well as tissue thickness and composition (e.g. distribution of symbionts and coral host pigments), and overall colony morphology. Hence the irradiance distribution between two morphologically similar coral fragments can vary, and in a reef environment a coral colony could experience varying external and internal light fields throughout the day, depending on the sun angle. Such heterogeneity in the light microenvironment across tissue and over the coral colony surface might present different optical niches that can drive phenotypic or genotypic diversification of symbionts (Lichtenberg et al., 2016) with different light adaptation and bleaching resistance across the same colony or between colonies with different morphology. It is, however, difficult to account for such spatial heterogeneity with microscale light measurements, especially for intra-tissue measurements that often rely on making a small incision in the coral tissue for probe insertion (Wangpraseurt et al., 2012). We show here that simulation of the spatial light distribution in corals with a known morphology and tissue structure is a powerful supplement to fine scale measurements of coral light fields.

### 3.2 Oxygen microenvironment

The dissolved O_2_ distribution in and around a coral colony for a given light field or in darkness is affected by the orientation of the coral with respect to the water flow field. Thus, different parts of the colony will experience changes in the thickness of the MBL and DBL depending on their flow exposure, which will translate into differences in O_2_ concentration within and at the surface of tissue.

The 3D simulations of light-driven O_2_ production in the coral fragment within the flow-cell were executed under two fragment orientations with respect to the water flow: with the groove between the two branches shaded against the flow by one branch (Figure 3a), and with the groove and both branches directly exposed to the flow (Figure 3e). The simulations showed higher O_2_ concentration in parts of the fragment that were shaded from the flow and exhibited a thicker DBL, as compared to more exposed areas (Figure 4a, d). The O_2_ concentration was high at the skeleton/tissue interface, due to the low diffusivity of the skeleton matrix (Figure 4b, c, e, and f).

**Figure 3:**
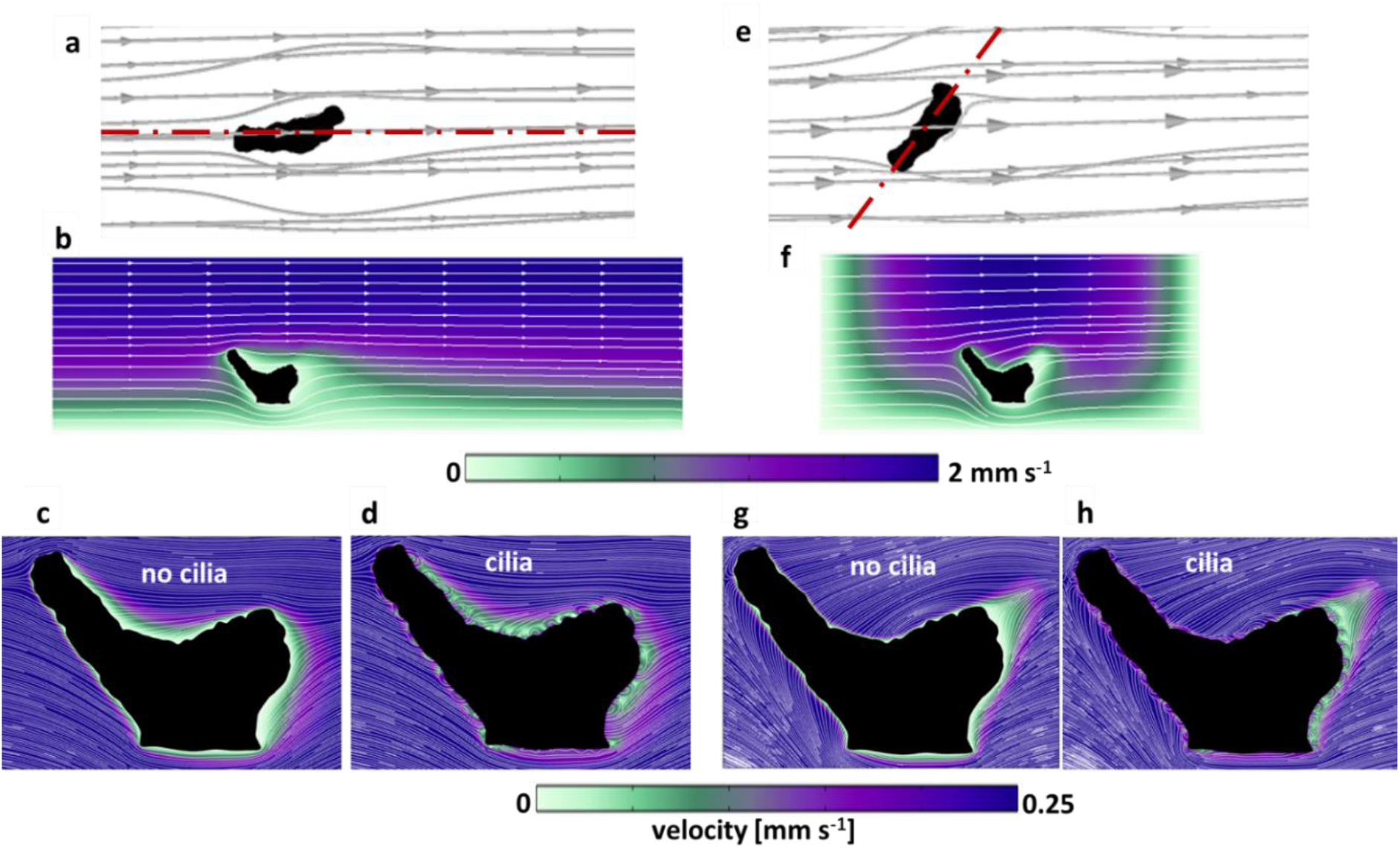
Computed flow field around the coral fragments in two orientations relative to the flow. a,b,c,d: aligned with the flow; e,f,g,h: across the flow. a,e: 3D streamlines of water flowing around the coral fragment. b,f: flow velocity magnitude (color map) and streamlines in the planar sections indicated in (a) and (e) with red dash-dot lines. c,d,g,h: details of velocity magnitude and streamlines in the neighborhood of the coral surface, with and without ciliary movement (same section planes as in b and f, respectively).

**Figure 4:**
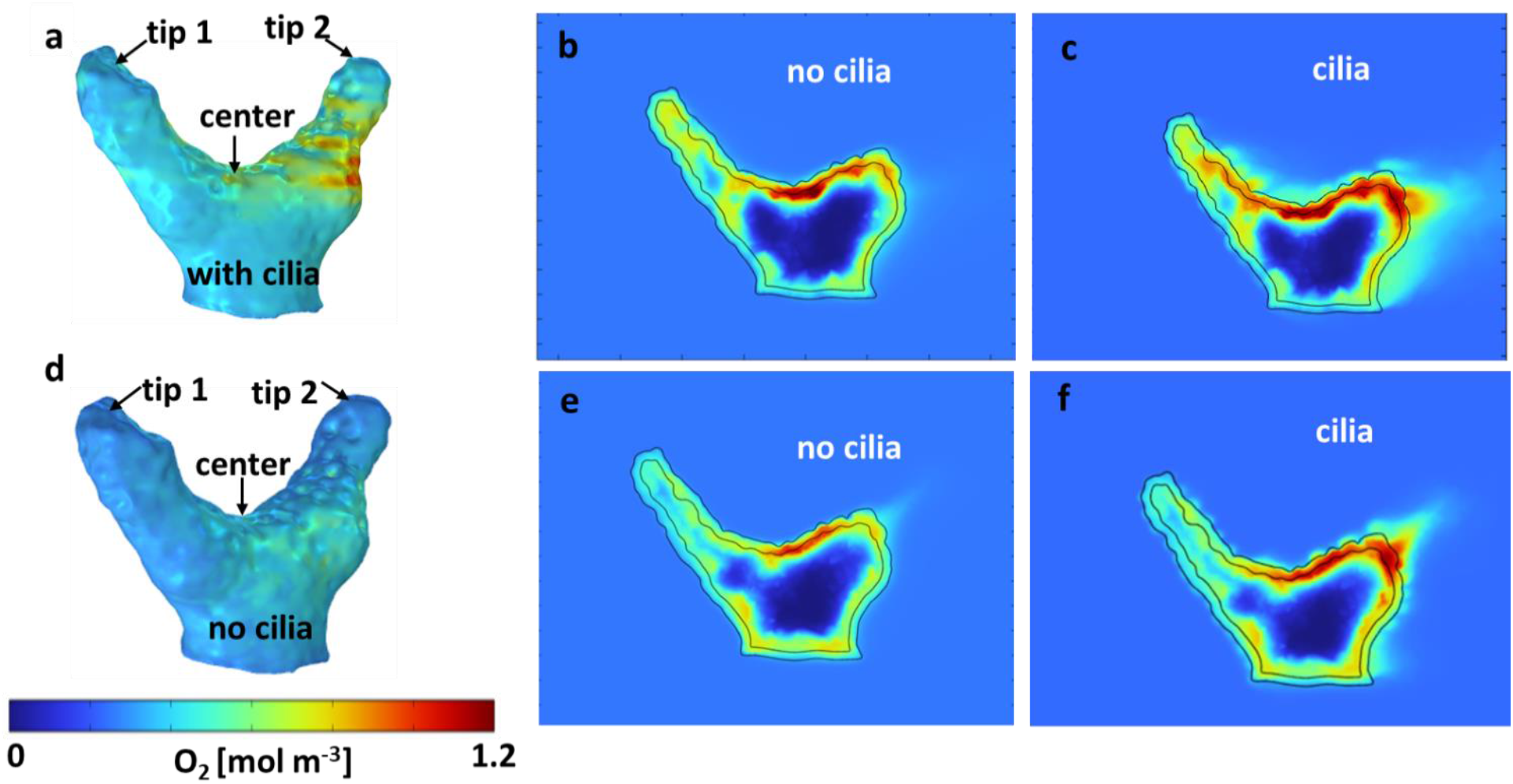
Computed dissolved O_2_ concentration in and around the coral fragment. a,b,c: coral fragment aligned with the flow; d,e,f: coral fragment across the flow. a,d: 3D O_2_ distribution on the coral (tissue/water) surface. b,e: 2D cross-sections of O_2_ concentration without ciliary movement. c,f: 2D cross-sections of oxygen concentration with ciliary movement. The flow direction and 2D sectioning planes are indicated in Figure 3a and 3e.

When the coral fragment was oriented with one branch facing the flow (named aligned with the flow in Figure 3a), the groove and the second branch exhibited a thicker DBL (Figure 3b), as compared to the orientation across the flow (Figure 3e, 3f) where the groove of the fragment was more flow-exposed. By constructing iso-velocity lines (IVL) at, for example, 1 mm s^-1^, the hydrodynamic boundary layer, in the more closed-groove orientation appears thicker (Figure S8a). This indicates a retarded water flow, as compared to a more open-groove orientation (Figure S8b) that showed IVL more conformal and closer to the coral surface. Vertical O_2_ concentration profiles extracted from the simulated O_2_ distributions (Figure S5) were compared with O_2_ microsensor measurements done within similar areas of the fragment, and showed a fairly good match (Figure 5) for the two different orientations of the fragment with respect to the water flow.

Under low or no flow conditions, mass transfer and therefore O_2_ concentration in corals can also be affected by the movement of cilia covering the coral ectoderm, which can create vortices and some advective transport at the coral tissue surface (Pacherres et al., 2020; Pacherres, 2022). We investigated the role of the ciliated coral tissue surface by comparing O_2_ microenvironment simulations with and without ciliary beating (Figure 4b, c). When no cilia beating was included, the simulations indicate a pronounced O_2_ accumulation at the center of the fragment (Figure 4b).

**Figure 5:**
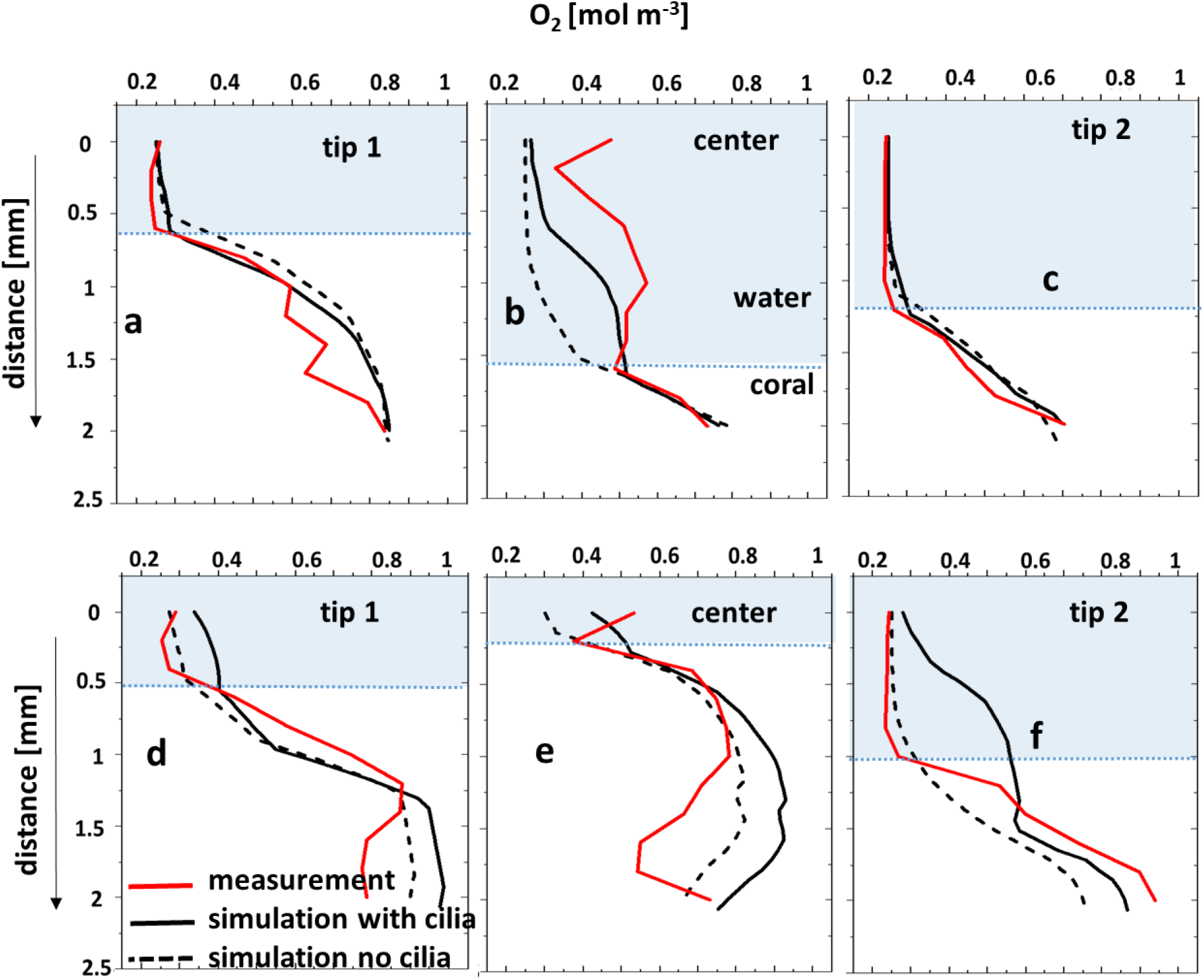
Comparison between microsensor-measured O_2_ profiles and the corresponding simulated profiles through the water column and tissue, with and without considering the surface ciliary motion. Coral fragment, a,b,c: aligned with the flow; d,e,f: across the flow. “center”, “tip 1” and “tip 2” measurement positions are indicated in Figure 4. The tissue/water boundary indicated here was determined from the simulated DO profile and compared with the measurements.

Whereas, when cilia movement was included, the simulated O_2_ distribution was more homogenous due to enhanced advective transport by the ciliary vortex formation (Figure 4c), in line with published experimental data (Pacherres et al., 2020). The comparison between simulated O_2_ profiles and the corresponding measurements (Figure 5a-c) also showed a closer match when cilia beating was included in the model. However, when the coral fragment was oriented across the flow, the central area was more exposed and the MBL was relatively thin.

Consequently, the measured and simulated O_2_ concentration profiles indicated a smaller impact of the ciliary movements on mass transfer (Figure 4d-f, 5d-f). This suggests that cilia-induced advective transport has a more significant effect on the O_2_ transport between the coral tissue and the surrounding seawater in colony regions with slow flow. We note that movements in the coral such as tissue-contraction and expansion and tentacle movement, which are presently not included in the model, could also potentially influence the O_2_ mass transport (Malul et al., 2020; Patterson, 1992).

### 3.3 Temperature microenvironment

Heat transfer simulations and temperature measurements were executed for both orientations of the coral sample with respect to the flow (Figure 6 - aligned; Figure S9 - across). The simulated temperature profiles with and without ciliary movement were very similar, showing an increase of less than 0.2°C, as visible in the 2D cross-sectional images of the computed temperature distribution (Figure 6b,c and Figure S9b,c) and extracted profiles (Figure 6d-f and Figure S9d-f). Temperature profiles measured at three different locations (Figure 6a, S9a) in the fragment were within the range of simulated temperatures, but with considerable noise. The water flow affected local heat transfer, where the temperature simulations show higher tissue heating from the absorbed light in fragment regions exposed to slower flow, albeit the absolute temperature increase was very moderate for the investigated branched coral.

**Figure 6:**
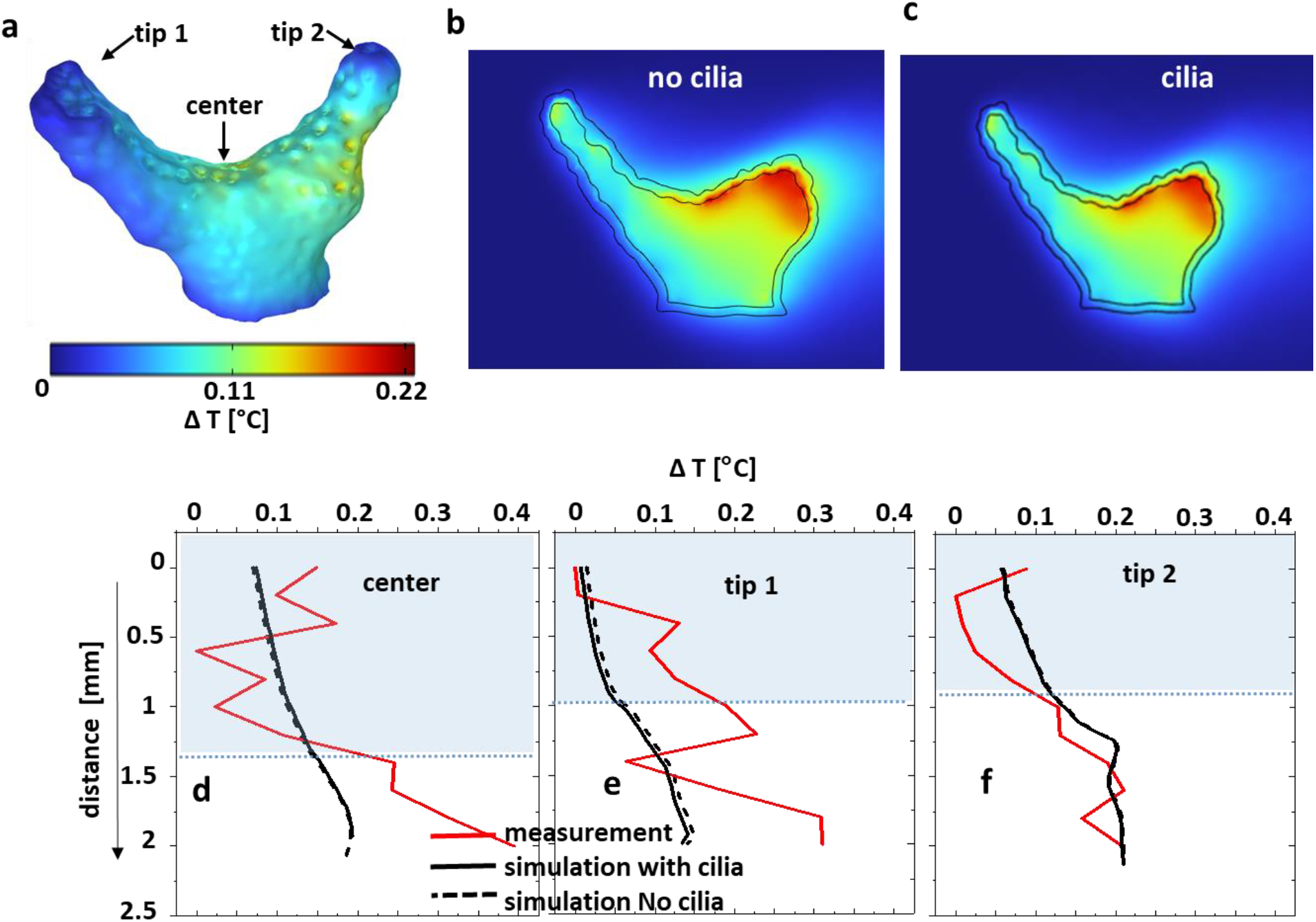
Comparison between measured and simulated temperature distribution for the coral fragment aligned with the flow. The temperature differences Δ*T* are between the local values and the inflow temperature. a: computed 3D Δ*T* on the coral (tissue/water) surface; b,c: Δ*T* in 2D cross-sections through the coral without and with ciliary movement. Sectioning planes are indicated in Figure 3a. d,e,f: measured and simulated temperature difference profiles through the water column and tissue, with and without considering the surface ciliary motion.

Simulations indicate that the slower heat transfer (thicker MBL and TBL) over tissue regions less exposed to flow was not significantly enhanced by ciliary movement. The temperature increase was highest near the skeleton/tissue interface, due to heat-insulating properties of the skeleton (Jimenez et al., 2008).

Clearly, it is difficult to make a systematic comparison between simulated and measured temperature profiles in this case (Figure 6d-f and Figure S9d-f) due to the very small temperature gradient between coral tissue and water close to the noise level of the microsensor measurements. The small temperature differences in the branched coral fragment are due to a high surface area to volume ratio (Jimenez et al., 2008), allowing a quick heat dissipation into the environment. There is thus a need for more accurate temperature microsensors, but it could also be relevant to compare simulated and measured temperatures on massive corals that have been shown to exhibit a stronger heating than branched corals (Jimenez et al., 2008; Jimenez et al., 2011; Jimenez et al., 2012). Last but not least, there is a lack of thermal property measurements in the different layers in coral tissue, and our simulation of the temperature microenvironment thus largely relied on values for tissue thermal properties from the biomedical literature.

### 3.4 Outlook

We developed a multiphysics modelling approach that enables simulation of the physico-chemical microenvironment of corals with a known 3D structure under defined irradiance and laminar flow conditions similar to commonly used flow chamber experimental setups. The model results were evaluated on 3D scanned coral samples, previously characterized with microsensor measurements of light, temperature and O_2_ in a laminar flow chamber. Such comparison generally showed a good agreement between measured and simulated data, but several model improvements can be considered for future work. The current model only considers light of individual wavelengths at a time and does not include inelastic scattering (e.g. conversion of light energy due to fluorescence). Broadband spectral simulations would e.g. enable simulations of how the spectral red shifts caused by fluorescent host pigments (Alieva et al., 2008; Salih et al., 2000), wavelength-specific reflectivity by chromoproteins (Alieva et al., 2008; D’Angelo et al., 2008) affect the coral light field and photosynthesis (Ben-Zvi et al., 2021), internal heat generation (Quick et al., 2018), as well as coral bleaching and recovery (Grinblat et al., 2018). However, an important prerequisite for more detailed simulations would be a better quantification of the inherent optical and thermal properties of coral tissue and skeleton, which are still very scarce in the literature. The present models therefore partially rely on assumed thermal property values taken from biomedical tissue studies.

The simple laminar flow scenario currently implemented in the present study does not cover all flow scenarios of corals *in situ*, and is more representative of calm sea conditions on reef flats and inside coral patches (Jimenez et al., 2011; Jimenez et al., 2012), as well as in many experimental flow chambers used for ecophysiological studies of coral metabolism. It is possible to include more complex flow scenarios in our modeling approach such as turbulent and oscillating flows (Ong et al., 2012; Ong et al., 2012), albeit at higher computational costs. Furthermore, it would be interesting to compare simulated and measured flow fields around corals, e.g. using particle imaging velocimetry (Pacherres et al., 2020) or combined measurements of flow and O_2_ fields around corals (Ahmerkamp et al., 2022; Pacherres et al., 2022). The latter could also enable implementation of a more detailed account for the role of ciliary movement for coral mass and heat transfer.

The present model only accounts for photosynthetic O_2_ production of symbionts using a simple approach based on reported quantum efficiencies, while O_2_ consumption by respiration in different tissue layers is partially based on published values and P/R relationships. The model could be expanded to better account for photosynthetic light saturation and photoinhibition.

Considering other chemical species (inorganic and organic carbon, various acid/base couples) and reactions in our model would enable the representation of calcification and carbon transfer between symbionts and host, and such work is in progress. Another desired expansion could be a more detailed account of metabolic processes in the coral skeleton, performed by the endolithic algae and microbes (Ricci et al., 2019; Tandon et al., 2022).

Furthermore, the present model assumes homogenous tissue thickness over the coral colony and does not include an accurate representation of fine scale topographic and anatomic features of coral tissue and skeleton, such as the polyps with tentacles and a complex internal gastrovascular system, and the more homogenous connective tissue between polyps. Our model also represents coral tissue as a static structure. This is a simplification, as corals exhibit pronounced tissue plasticity via contraction and expansion, which can strongly affect surface area to volume ratios (Patterson, 1992) and optical properties (Wangpraseurt et al., 2017), which in turn modulate mass, heat and radiative transfer processes. More precise tomographic mapping of coral tissue and skeleton morphology and thickness e.g. with μCT scanning or OCT, in combination with including fluid-structure interaction models with moving interfaces (Taherzadeh et al., 2012) could allow for simulating effects of coral topography and mechanics on the coral microenvironment, but this will require substantial computational resources.

Finally, the presented 3D modelling approach can also be used to simulate different time-dependent environmental conditions (e.g., variable flow, day-night hypoxia, different solar irradiation regimes), which can help evaluating mechanisms driving coral stress responses as well as basic niche shaping factors for symbionts and microbiomes in the coral holobiont. The model could also be useful in more applied research such as in the ongoing attempts to create bionic corals (Wangpraseurt et al., 2022; Wangpraseurt et al., 2020) or for optimization of other 3D bioprinted constructs (Krujatz et al., 2022), where different designs can be evaluated and optimized. Our approach is also relevant for simulating structure-function relationships in other benthic systems such as photosynthetic biofilms and aquatic plant tissue, and can also be adapted to other sessile organisms such as symbiont-bearing giant clams, ascidians, jellyfish or foraminifera.

## 4. Conclusion

We developed a 3D multiphysics model to simulate the spatial distribution of light, O_2_ and temperature (and the corresponding radiative, mass and heat transfer) across a stratified coral tissue, skeleton and around natural coral morphologies exposed to a defined flow field and incident irradiance. Model results compared well with spatial measurements of light, O_2_ and temperature under similar flow and light conditions. The model reveals how the interaction between incident irradiance, water flow and complex coral morphology leads to pronounced spatial heterogeneity and microenvironments, both across tissue layers and between different areas of the coral. Such simulation of the physico-chemical microenvironmental landscape using real 3D scanned coral structures as an input, can i) give insights to coral tissue and skeleton compartments that are difficult to reach with existing sensor technology, and ii) identify hotspots of activity that can inform detailed measurements e.g. with microsensors and/or various bioimaging techniques for mapping structure and function. The model can simulate effects of different environmental (e.g. light, temperature, hypoxia or flow) and structural (e.g. symbiont and host pigment density and distribution, and the distribution of endolithic microbes) factors on the coral microenvironment and metabolic activity, which can be tested experimentally. We argue that such combination of modeling and experimental investigation is a strong tool set for unravelling structure-function relations and basic regulatory mechanisms in coral biology and stress responses, including effects of climate change and other anthropogenic threats.

## Acknowledgements

We acknowledge technical assistance with coral husbandry by Sofie Jakobsen, Victoria Thuesen and Caroline Vigsbo Christensen. This study was funded by the Gordon and Betty Moore Foundation through grant no. GBMF9206 to M.K. (https://doi.org/10.37807/GBMF9206). C.P. acknowledges support and access to computational resources offered by the KAUST Computing Center.

## Author Contributions

S.M., C.P. and M.K. designed the research. S.M. developed the coral model, performed computer simulations and measurements. S.M., C.P. and M.K. analyzed the data. S.M., C.P. and M.K. wrote the manuscript. C.P. and M.K. provided research infrastructure.

## Data availability

All data will be made available in Dyrad, and scripts of the modeling approaches are available from the authors at reasonable request.

