## Supplemental Table 1 for "Modeling the radiative, thermal and chemical microenvironment of 3D scanned corals"

### Supplementary information

**Table S1. Model parameters**

| Parameter | Symbol | Value | Unit | Source |
|---|---|---|---|---|
| **Geometry** | | | | |
| Length, flow chamber | $L_x$ | 190 | mm | Measured |
| Width, flow chamber | $L_y$ | 55 | mm | Measured |
| Height, flow chamber | $L_z$ | 80 | mm | Measured |
| Coral skeleton | | | mm | 3D scan leading to an STL file |
| Coral layers | | | | |
| mucus | $L_{muc}$ | 0.08 | mm | (Lichtenberg et al., 2016) |
| epidermis | $L_{ep}$ | 0.08 | mm | (Lyndby et al., 2016) |
| mesoglea | $L_{mes}$ | 0.08 | mm | (Lichtenberg et al., 2016) |
| oral gastrodermis | $L_{og}$ | 0.1 | mm | (Lichtenberg et al., 2016) |
| gastrovascular cavity | $L_{gas}$ | 0.08 | mm | chosen |
| aboral gastrodermis | $L_{ag}$ | 0.1 | mm | (Lichtenberg et al., 2016) |
| total tissue thickness | $L_{tis}$ | 0.65 | mm | Mean of OCT and 3D scans |
| **Optics** | | | | |
| Absorption coefficients | | | | |
| Water | $\mu_{a,wat}$ | $3.6 \cdot 10^{-3}$ | $cm^{-1}$ | (Kirk, 1994) |
| Tissue | $\mu_{a,tis}$ | 1.8 | $cm^{-1}$ | (Wangpraseurt et al., 2016) |

| | | | | |
|---|---|---|---|---|
| | | | | (Jacques et al., 2019) |
| Skeleton | $\mu_{a,skel}$ | 0.01 | $cm^{-1}$ | (Wangpraseurt et al., 2016) (Jacques et al., 2019) |
| Scattering coefficients | | | | |
| Water | $\mu_{s,wat}$ | $1 \cdot 10^{-5}$ | $cm^{-1}$ | (Kirk, 1994) |
| Tissue | $\mu_{s,tis}$ | 100 | $cm^{-1}$ | (Wangpraseurt et al., 2016) (Jacques et al., 2019) |
| Skeleton | $\mu_{s,skel}$ | 34 | $cm^{-1}$ | (Wangpraseurt et al., 2016) (Jacques et al., 2019) |
| Anisotropy coefficients | | | | |
| water | $g_{wat}$ | 1 | - | (Kirk, 1994) |
| tissue | $g_{tis}$ | 0.9 | - | (Wangpraseurt et al., 2016) (Jacques et al., 2019) |
| Skeleton | $g_{skel}$ | 0.9 | - | (Wangpraseurt et al., 2016) (Jacques et al., 2019) |
| Refractive indices | | | | |
| water | $n_{wat}$ | 1.33 | - | (Kirk, 1994) |
| Tissue | $n_{ag}$ | 1.38 | - | (Wangpraseurt et al., 2016) |
| Skeleton | $n_{skel}$ | 1.66 | - | (Ghosh, 1999) |
| **Flow and heat** | | | | |
| maximum ciliate beating velocity | $C_{vel}$ | 150 | µm $s^{-1}$ | (Shapiro et al., 2014) |

| | | | | |
|---|---|---|---|---|
| characteristic length scale of the vortices | $\delta$ | 1200 | µm $s^{-1}$ | OCT scans |
| Viscosity water | $\eta_{wat}$ | 0.0009 | Pa s | (DeWitt, 1990) |
| Densities | | | | |
| Water | $\rho_{wat}$ | 997 | kg $m^{-3}$ | (DeWitt, 1990) |
| tissue (all) | $\rho_{tis}$ | 1109 | kg $m^{-3}$ | (Hasgall PA, 2018) |
| Skeleton | $\rho_{skel}$ | 2930 | kg $m^{-3}$ | (Bragg, 1924) |
| Heat conductivities | | | | |
| Water | $k_{wat}$ | 0.6 | W $m^{-1}$ $K^{-1}$ | (DeWitt, 1990) |
| tissue (all) | $k_{tis}$ | 0.037 | W $m^{-1}$ $K^{-1}$ | (Hasgall PA, 2018) |
| Skeleton | $k_{skel}$ | 0.5 | W $m^{-1}$ $K^{-1}$ | (Jimenez et al., 2008) |
| Heat capacities | | | | |
| Water | $C_{p,wat}$ | 4183 | J $kg^{-1}$ $K^{-1}$ | (DeWitt, 1990) |
| tissue (all) | $C_{p,tis}$ | 300 | J $kg^{-1}$ $K^{-1}$ | (Hasgall PA, 2018) |
| Skeleton | $C_{p,skel}$ | 1846 | J $kg^{-1}$ $K^{-1}$ | (Jimenez et al., 2008) |
| **Mass transport** | | | | |
| Diffusion coefficients | | | | |
| $O_2$ in water | $D_{O2,wat}$ | $2 \cdot 10^{-9}$ | $m^2$ $s^{-1}$ | (Haynes, 2016) |
| $O_2$ in tissue (all) | $D_{O2,tis}$ | $1 \cdot 10^{-9}$ | $m^2$ $s^{-1}$ | (MacDougall & McCabe, 1967) |

| | | | | |
|---|---|---|---|---|
| $O_2$ in skeleton | $D_{O2,skel}$ | $2 \cdot 10^{-11}$ | $m^2\ s^{-1}$ | From water, assumed 1% porosity |
| Gas-liquid mass transfer coefficient | $k_l a$ | 10 | $s^{-1}$ | Value adjusted to avoid excessive $O_2$ accumulation |
| Oxygen solubility in water | $c_{s,O2}$ | 1.25 | $mol\ m^{-3}$ | (Xing et al., 2014) |
| Quantum efficiency oral gastrodermis | $Q_{og}$ | 0.02 | $mol\ mol^{-1}$ | (Brodersen et al., 2014) |
| Quantum efficiency aboral gastrodermis | $Q_{ag}$ | 0.01 | $mol\ mol^{-1}$ | (Brodersen et al., 2014) |
| P/R ratio in gastrodermis | $PR$ | 3.5 | - | (Cooper et al., 2011) |
| $O_2$ half saturation coefficient | $K_{O2}$ | $3.13 \cdot 10^{-3}$ | $mol\ m^{-3}$ | (Bowie et al., 1985) |
| Maximum $O_2$ consumption rate in the skeleton | $R_{cp,skel}$ | $3.75 \cdot 10^{-5}$ | $mol\ m^{-3}\ s^{-1}$ | (Kühl et al., 2008) |
| Average water velocity | $v_0$ | 0.1 | $cm\ s^{-1}$ | measurement |
| Ambient temperature | $T_0$ | 26 | °C | measurement |
| Inlet $O_2$ concentration | $c_{0,O2}$ | 0.25 | $mol\ m^{-3}$ | measurement |

| | | | | |
|---|---|---|---|---|
| Incident downwelling photon irradiance | $I_0$ | 1300 | µmol photon $m^{-2}$ $s^{-1}$ | measurement |

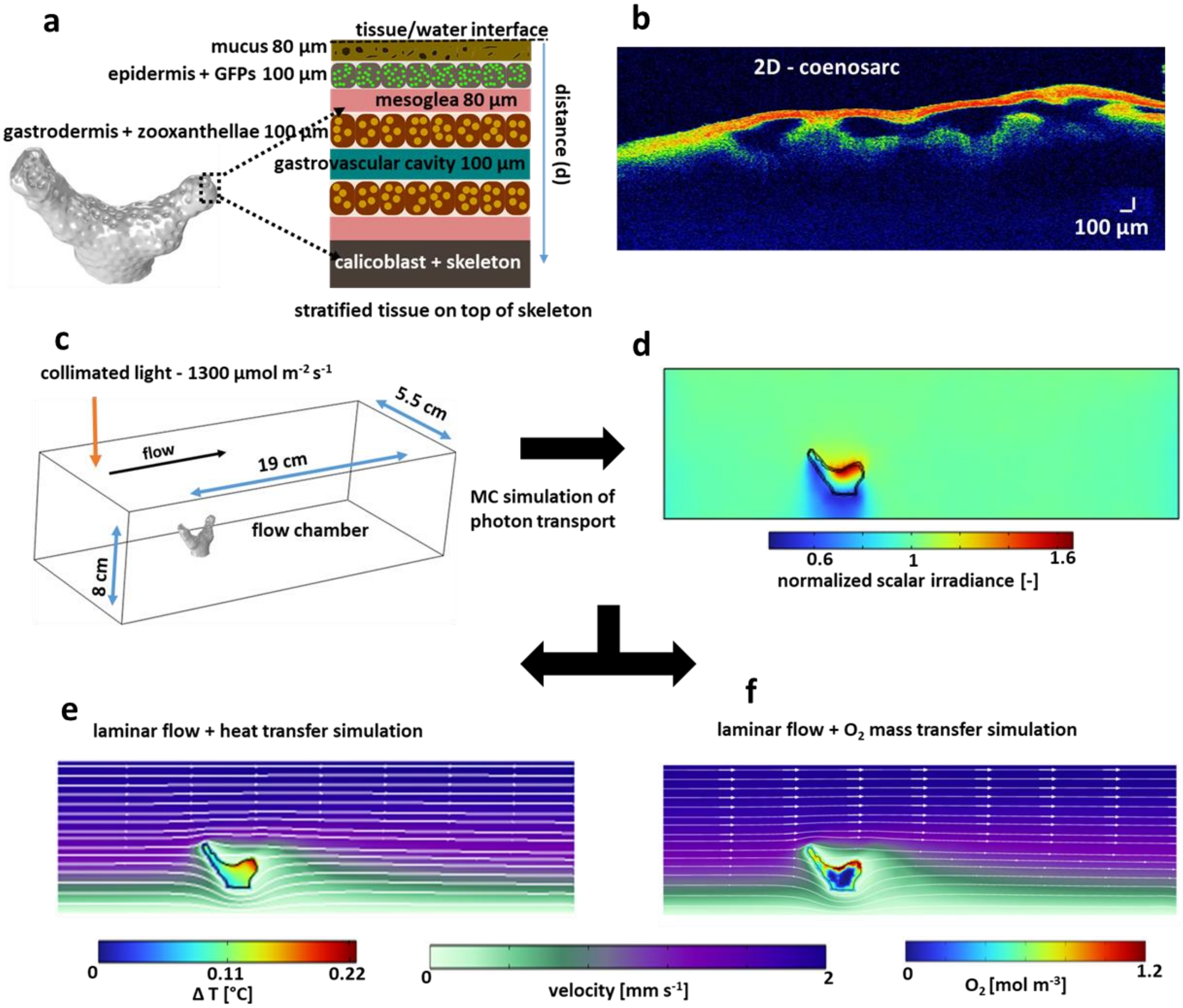


Figure S1: Schematic of the steps involved in the simulation. a: Insert showing stratified tissue covering the skeleton; b: 2D OCT image of coenosarc; c: 3D model geometry consisting of the coral fragment in a flow chamber, used for simulation; d: Simulated scalar irradiance along the middle plane; e: Simulated laminar flow and heat distribution along the middle plane; f: Simulated laminar flow and $O_2$ distribution along the middle plane. Sectioning shown in Figure 3a.

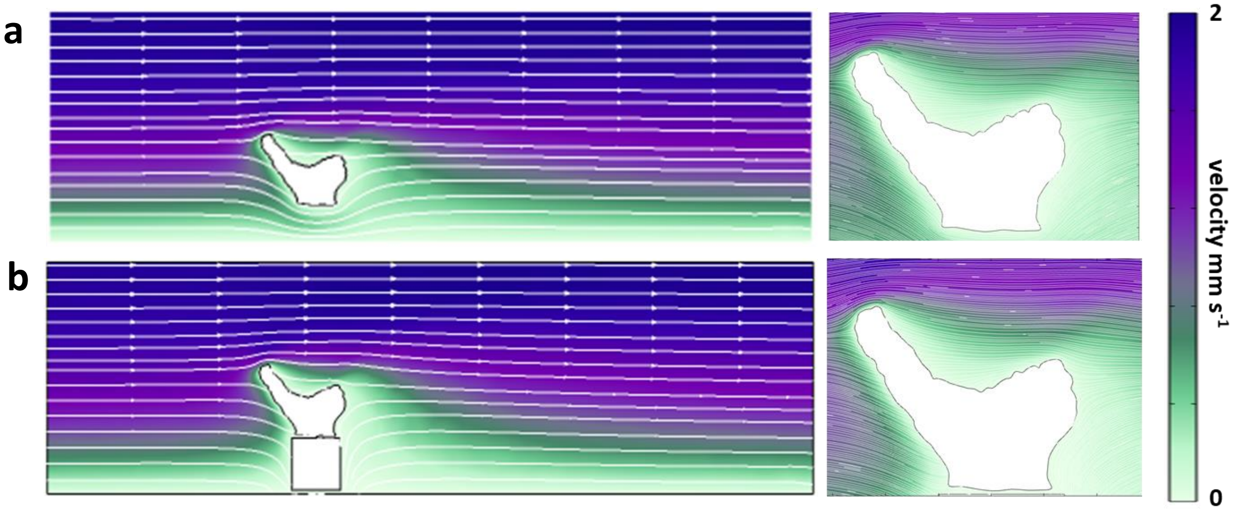


Figure S2: Comparison of flow simulation around the fragment with (a) and without (b) inclusion of the base.

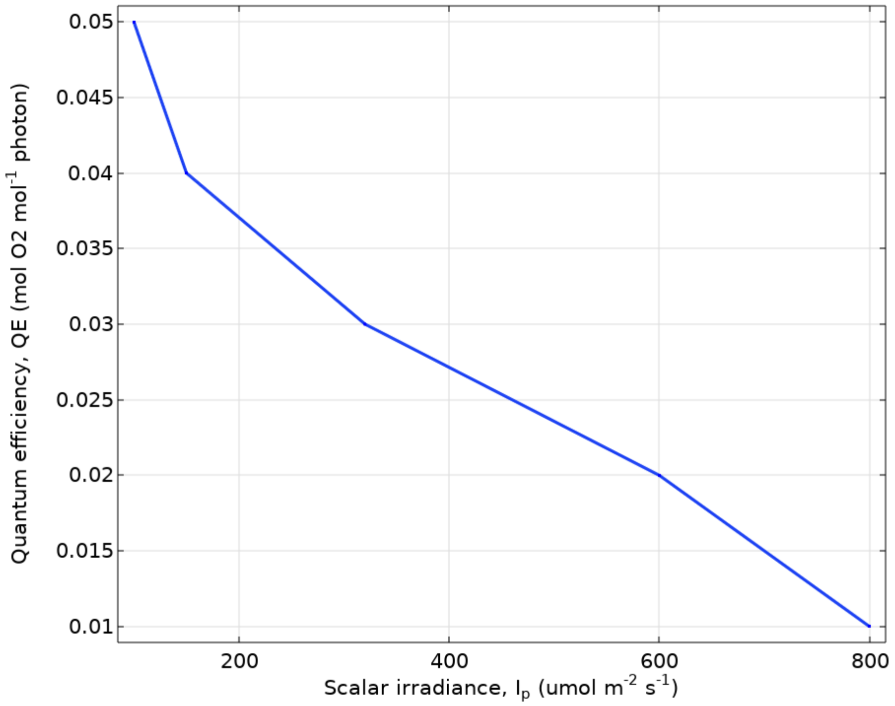


Figure S3: Quantum efficiency of photosynthesis, *QE*, as a function of scalar irradiance, $I_p$. (Brodersen et al., 2014)

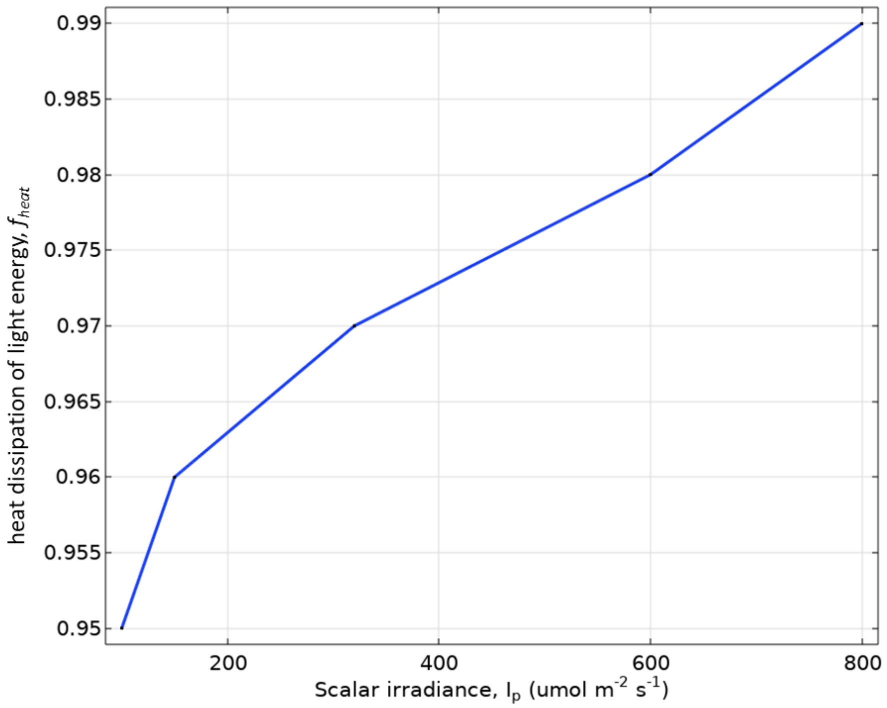


Figure S4: Fraction of light energy dissipated as heat $f_{heat}$, as function of scalar irradiance $I_p$. (Brodersen et al., 2014)

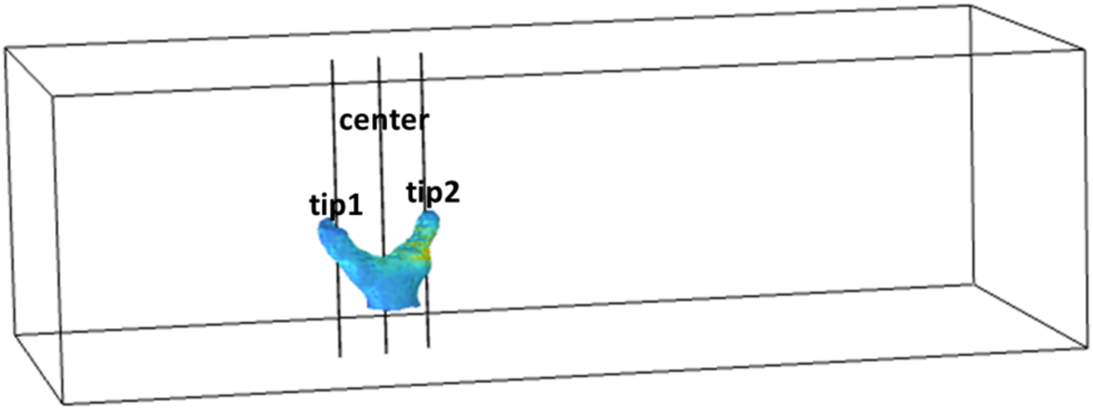


Figure S5: Sketch indicating the vertical line profiles extracted from the simulation data.

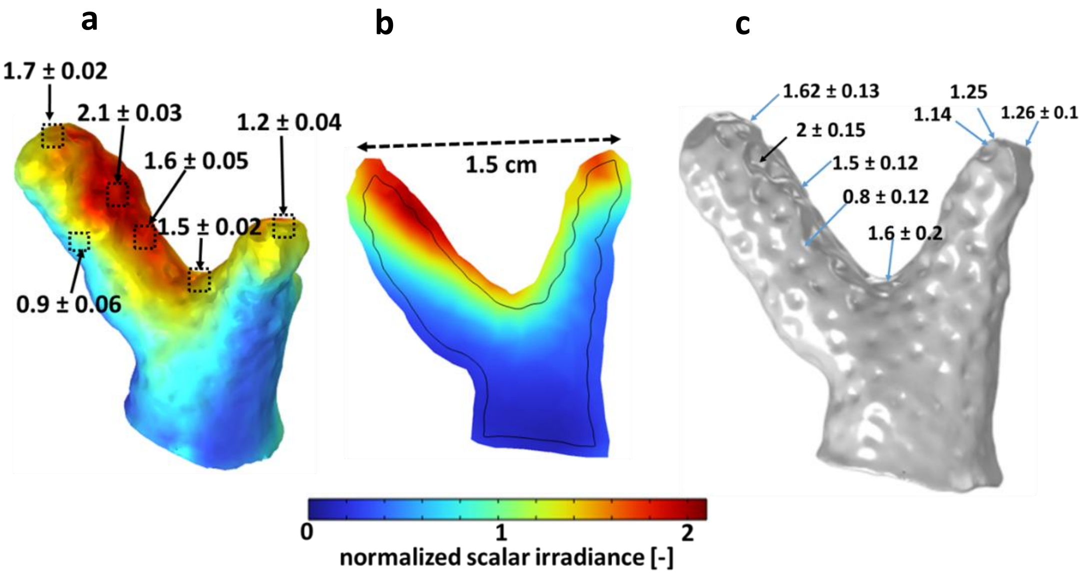


Figure S6: Comparison between measured and simulated scalar irradiance normalized to the incident irradiance at 636 nm for fragment 2. a: Simulated scalar irradiance at the tissue water interface; b: Cross-section of simulated scalar irradiance along the center of the fragment; c: Measured scalar irradiance at 636 nm. Tissue optical properties (OP2)(Wangpraseurt et al., 2018): $\mu_a = 1.8\ cm^{-1}$, $\mu_s = 40\ cm^{-1}$, g = 0.51, n=1.38. Skeleton optical properties: $\mu_a = 0.01\ cm^{-1}$, $\mu_s = 40\ cm^{-1}$, g = 0.28, n = 1.66. The values shown here are mean ± standard deviation, averaged over 12 different points.

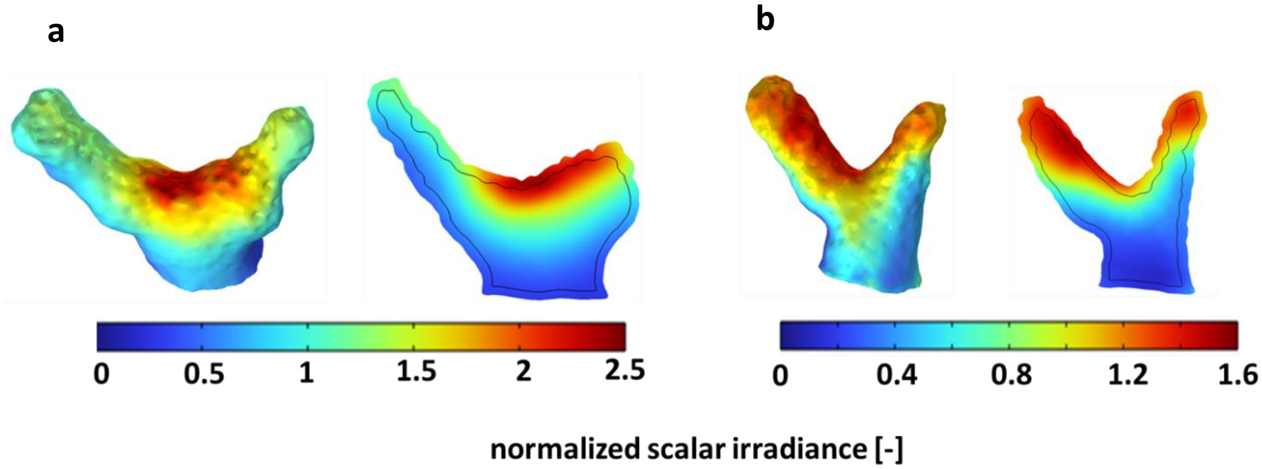


Figure S7: Simulated scalar irradiance for a: Fragment 1 assuming optical properties OP2. b: Fragment 2 assuming optical properties OP1 from Table 1.

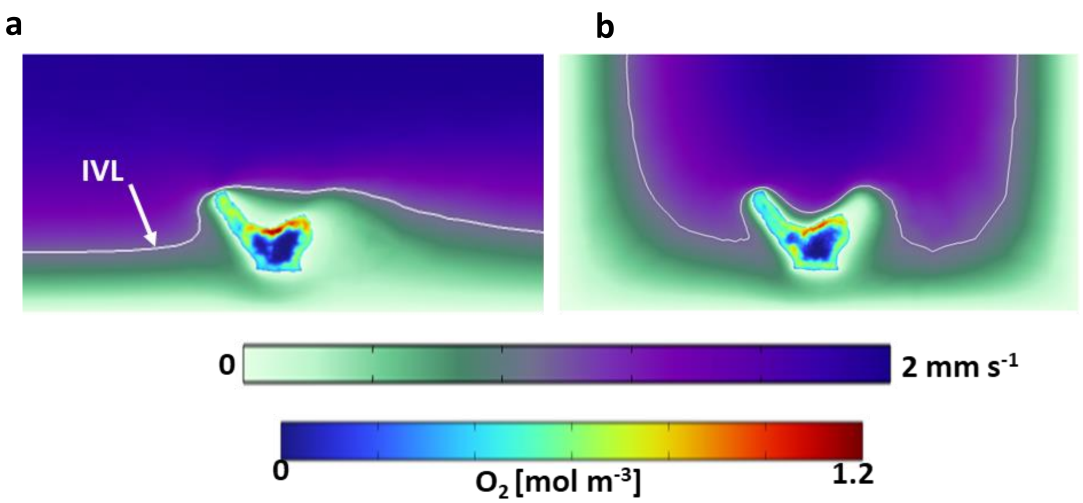


Figure S8: 2D cross-sectional view indicating the iso-velocity line for 1 mm $s^{-1}$ flow around the fragment and the simulated $O_2$ concentration in the coral fragment with the fragment a: aligned with flow (Figure 3a) and b: across the flow (Figure 3e).

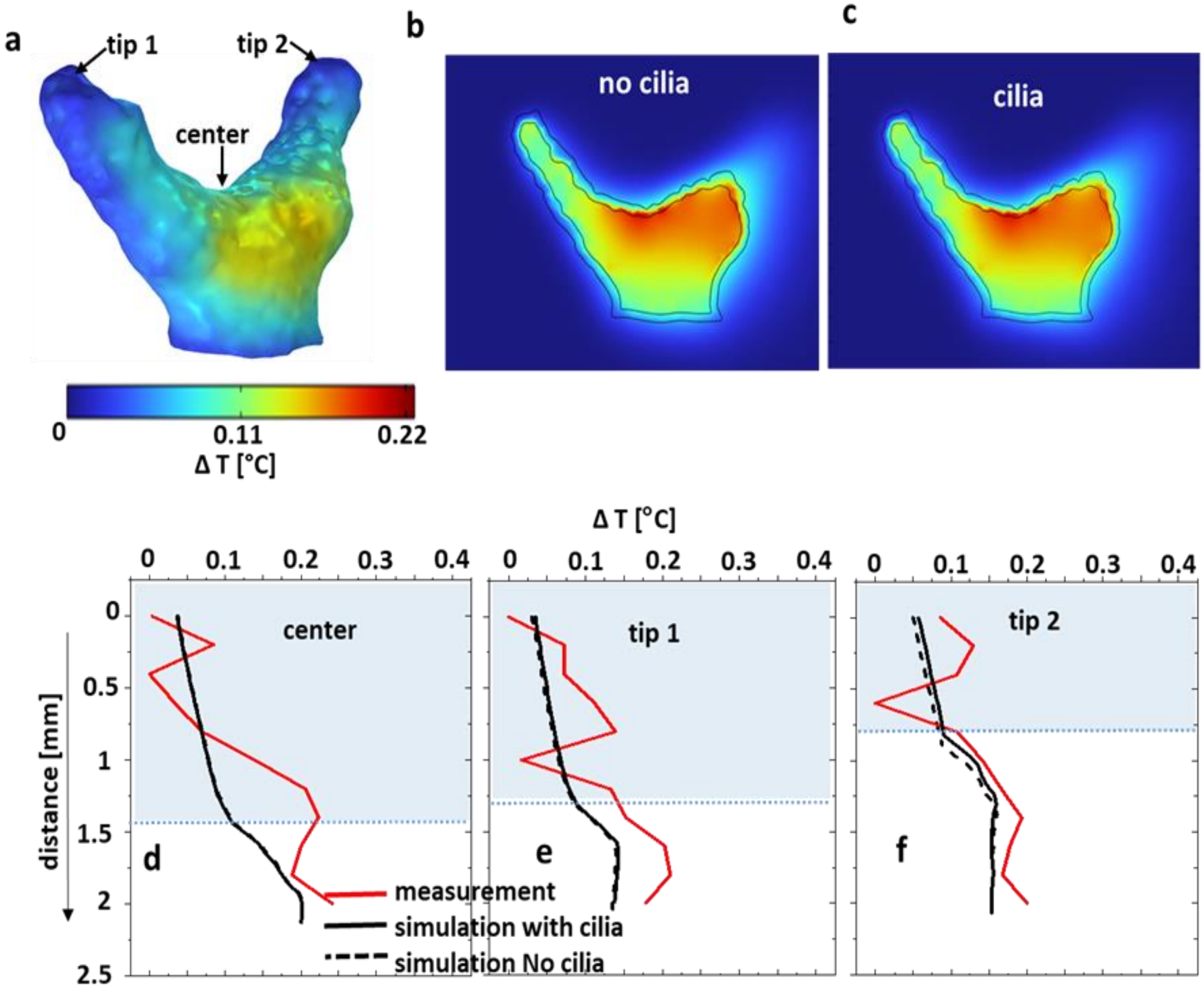


Figure S9: Comparison between measured and simulated temperature distribution for the coral fragment across the flow. The simulated temperature differences, $\Delta T$, are between the local values and the inflow temperature. a: computed $\Delta T$ over the coral tissue-water interface; b,c: $\Delta T$ in 2D cross-sections through the coral, simulated without and with ciliary movement. Sectioning planes are indicated in Figure 3e. d,e,f: measured and simulated temperature difference profiles through the water column and tissue, with and without considering the surface ciliary motion.

75
